# CD163⁺/Dab2⁺ Macrophages Alleviate Cardiac Hypertrophy via Nrg2/ErbB4-Mediated Mitochondrial Reprogramming

**DOI:** 10.1101/2025.05.22.655661

**Authors:** Qi Zhu, Shuilin Liao, Yin Zhou, Ruoyu Zhou, Ming Dong

## Abstract

**Background:** Pathological cardiac hypertrophy is a hallmark of numerous cardiovascular diseases, yet effective targeted therapies remain elusive in current clinical practice. Cardiac macrophages contribute to disease progression, but the underlying mechanisms have not been fully elucidated.

**Methods:** Single-nucleus RNA-sequencing, bulk RNA-sequencing, proteomics, metabolomics, and a mouse model of pressure overload were employed to investigate cardiac remodeling. We identified a macrophage subset co-expressing CD163 and Dab2 and examined its role using immunofluorescence, flow cytometry, and functional assays. We further assessed the Nrg2/ErbB4 signaling axis through genetic and pharmacological modulation.

**Results:** CD163⁺/Dab2⁺ macrophages were reduced in hypertrophic hearts and positively correlated with Nrg2 expression. These macrophages alleviated cardiomyocyte hypertrophy in vitro, an effect abolished by Nrg2 knockdown. In vivo, recombinant Nrg2 treatment mitigated cardiac hypertrophy, preserved mitochondrial structure, and restored bioenergetics via the ErbB4 receptor. Transcriptomic analyses confirmed enhanced expression of genes involved in mitochondrial oxidative phosphorylation. Furthermore, CD163^+^/Dab2^+^ macrophages improved mitochondrial dysfunction by the Nrg2/ErbB4 pathway in vitro.

**Conclusions:** We identified a CD163⁺/Dab2⁺ macrophage subset that protects against pathological cardiac hypertrophy by promoting mitochondrial function through Nrg2/ErbB4 signaling. This axis may offer a promising therapeutic target for interventions in pathological cardiac hypertrophy.

## Introduction

Cardiac hypertrophy initially serves as an adaptive response to hemodynamic stress in conditions such as hypertension or valvular disease^1^. However, sustained pathological stimuli induce detrimental myocardial remodeling characterized by irreversible cardiomyocyte hypertrophy, fibrotic deposition, inflammatory cell infiltration, and mitochondrial impairment, which ultimately culminate in heart failure^2^. Inhibiting hypertrophy represents a promising therapeutic strategy in heart failure, yet effective targeted treatments are still absent in current daily clinical practice, making it imperative to develop new therapies to counteract pathological cardiac hypertrophy^3^, particularly by elucidating the cellular and molecular mechanisms that drive its progression.

Recent studies have emphasized the regulatory roles of immune cells, particularly heterogeneous macrophage populations, in orchestrating cardiac remodeling through tissue repair, inflammatory modulation, and metabolic reprogramming^4,5^. Macrophages exhibit remarkable plasticity, dynamically polarizing into distinct subsets with divergent functions, which either exacerbate or ameliorate cardiac pathology depending on their activation state and microenvironmental cues^6^. Cardiac macrophages are broadly categorized into resident CCR2^-^ (CC-motif chemokine receptor 2) macrophages and recruited CCR2⁺ macrophages, each contributing uniquely to disease progression. Recruited CCR2⁺ macrophages, often polarized to a pro-inflammatory M1-like phenotype, secrete interleukin-6 (IL-6), tumor necrosis factor-α (TNF-α), and other cytokines that promote cardiomyocyte hypertrophy, fibroblast activation, and extracellular matrix deposition^7^. For instance, NLRC5, a macrophage-specific pattern recognition receptor, interacts with HSPA8 to suppress IL-6 release, thereby attenuating STAT3-mediated cardiomyocyte hypertrophy and fibrosis in pressure-overloaded hearts^8^. Resident cardiac macrophages express marker genes such as Timd4, Lyve1, and M2-like markers Cd163 and Mrc1 (Cd206)^7^. Resident Timd4⁺ macrophages exhibit reparative properties by resolving inflammation and facilitating tissue repair^5,9^. CD206^+^ macrophages accumulate in infarcted and border zones following acute myocardial infarction (MI), where they facilitate tissue repair and attenuate maladaptive cardiac remodeling^10^.

Metabolic dysregulation, particularly mitochondrial dysfunction, is central to the pathogenesis of pathological cardiac hypertrophy and heart failure^11^. The heart, a high-energy-demand organ, relies on mitochondrial oxidative phosphorylation to sustain its contractile function. However, under pathological stress, such as pressure overload or metabolic disturbances, mitochondrial dysfunction disrupts energy equilibrium, leading to oxidative damage, metabolic reprogramming, and maladaptive remodeling^12^. Emerging evidence highlights that macrophages not only modulate inflammatory responses but also directly influence mitochondrial homeostasis, thereby shaping the metabolic and functional fate of cardiomyocytes^13^. Genetic depletion of the phagocytic receptor MerTK or resident macrophages in mice results in the impaired autophagy, accumulation of anomalous mitochondria in cardiomyocytes, metabolic alterations, and ventricular dysfunction^14^. CD163^+^ macrophages protect against pressure overload-induced left ventricular dysfunction and mitochondrial dysfunction via IL-10-dependent pathways^15^. Additionally, cardiac-resident CD163^+^RETNLA^+^ macrophages with high expression of TREM2, which scavenge dysfunctional mitochondria and suppress inflammation via TREM2-dependent pathways, are critical for mitigating sepsis-induced cardiomyopathy (SICM)^16^. Therefore, mitochondrial dysfunction is considered a direct therapeutic target for improving cardiac function^17^.

Despite increasing recognition of the role of immune cells in cardiac remodeling, the identity of specific macrophage subsets involved and the molecular pathways through which they influence cardiomyocyte metabolism remain poorly defined. In this study, we sought to investigate the cellular and molecular mechanisms by which macrophages contribute to pathological cardiac hypertrophy, with a particular focus on macrophage–cardiomyocyte communication. Using multi-omics approaches and a pressure-overload mouse model, we aimed to identify functionally distinct macrophage subsets and uncover their signaling interactions with cardiomyocytes that may reveal novel therapeutic targets for intervention.

## Methods

An expanded Methods section is available in the online-only Data Supplement.

### Mouse Studies

All animal procedures were approved by the Institutional Animal Care and Use Committee of Guangzhou Medical University (Guangzhou, China) and conformed to the Guide for the Care and Use of Laboratory Animals (8th edition, National Academies Press, 2011). Mice were maintained in the AAALAC-accredited Center for Experimental Animals at Guangzhou National Laboratory under standardized husbandry conditions. To minimize bias, group assignments were implemented through a randomized, double-blind protocol. 8 weeks old male mice were used for experiments as indicated. Surgical details for TAC procedures and treatment regimens are provided in the Supplemental Methods.

### Transverse aortic constriction and Nrg2/AG1478 treatment

All animal experiments were approved by the Animal Care Committee of Affiliated First Hospital of Guangzhou Medical University and were performed according to the institutional guidelines. TAC surgery was used to construct a model of cardiac pressure overload. Briefly, 6-week-old mice were anesthetized using 3% isoflurane and placed on a heated surgical plane. After alcohol disinfection, a mid-sternal thoracotomy was performed to expose the trachea and underlying thymus. The thymus was then carefully separated from the aortic arch. A 6-0 silk suture was placed between the innominate artery and the left common carotid artery and a 27 G constriction needle was placed parallel to the aortic arch. The suture was tied securely around both the needle and the aorta, creating a consistent constriction with an approximate diameter of 0.45 mm. After carefully removing the needle, the chest was closed using 5-0 sutures, and then the skin was closed with 3-0 sutures. The mice were monitored until recovery in a 37 ℃ heated cage. Sham surgery was performed to include all the steps described above except for the aortic ligation.

For the activation and blockade of the Nrg2 and ErbB4 pathway, 1 week after TAC surgery, mice were randomized and continuously treated with either vehicle or recombinant Nrg2 (produced by our lab) at a dosage of 500 μg/kg/d and inhibitor AG1478 (HY-13524, MCE) at a dosage of 10 mg/kg/d for 7 days. At the indicated timepoints after TAC surgery, mice were euthanized and the heart tissues were harvested to analyze pathological cardiac hypertrophy.

### Data Availability Statement

Data, analytical methods, and study materials will be made available to qualified researchers upon reasonable request, pending compliance with applicable legal and institutional regulations.

### Statistical analysis

The statistical analysis was conducted using unpaired two-tailed unpaired Student’s t-tests for two groups, and one-way or two-way ANOVA followed by Tukey’s post hoc test for multiple groups. All statistical analyses were performed with GraphPad Prism (version 9.0) and SPSS (Version 20.0). Data were presented as means ± SD from individual experiments. Statistical significance was denoted as \**P*<0.05; \*\**P*<0.01; \*\*\**P*<0.001.

## Results

### Multi-omics analysis of pathological cardiac hypertrophy

To delineate the molecular trajectory of heart failure development, we performed longitudinal multi-omics profiling (single-nucleus RNA sequencing, bulk RNA-sequencing, proteomics, and metabolomics) on murine hearts at 0, 2, and 8 weeks post transverse aortic constriction (TAC) surgery (Figure 1A). TAC-induced pressure overload caused cardiac dysfunction at 8 weeks post TAC surgery, as evidenced by echocardiography showing reduced left ventricular ejection fraction (LVEF) and fractional shortening (LVFS), alongside increased left ventricular mass, volume, internal diameter, and posterior wall thickness compared to those of sham controls Figure 1B, C; Figure S1A). Hematoxylin and eosin (H&E) staining and Masson staining of cardiac sections indicated that myocardial fibrosis was prominently increased (Figure 1D, E) and wheat germ agglutinin (WGA) staining revealed an enlarged cardiomyocyte cross-sectional area after TAC surgery (Figure S1B, C). Consistent with disease progression, the mRNA levels of the hypertrophic markers, atrial natriuretic peptide (Anp) and brain natriuretic peptide (Bnp) were substantially upregulated (Figure S1D). Following stringent quality control and clustering analysis, a total of 53,957 cells and 28 clusters with distinct expression features were identified by unbiased clustering of the cellular aggregate using the Seurat software suite^18^ (Figure 1F, G; Figure S2A). As anticipated, the expression levels of the hypertrophy-related genes, such as Nppa (also known as Anp), Nppb (also known as Bnp), and myosin heavy chain 7 (Myh7) were gradually increased during pathological cardiac hypertrophy (Figure S2B). Eight major cell types were defined, cardiomyocytes (TNNT2^+^Ttn^+^), fibroblasts (Col1a2^+^Col3a1^+^), endothelial cells (Pecam1^+^Egfl7^+^), smooth muscle cells (Rgs5^+^Pdgfrb^+^), macrophages (Adgre1^+^Rbm47^+^), B cells (Cd22^+^Btla^+^), T cells (Skap1^+^Lef1^+^), and neuronal cells (Syt1^+^Nrxn3^+^), based on their respective molecular signatures (Figure 1H), and cell clusters were visualized in t-SNE dimensionality reduction plots (Figure 1I, J). A functional comparison of biological processes revealed the relationships among different cells and physiological activities, such as the specific enrichment of heart process and heart contraction genes in cardiomyocytes, the expression of mainly extracellular structure organization genes in fibroblasts, and the correlation between macrophages and myeloid leukocyte differentiation (Figure S2C). Single-nucleus transcriptional profiling uncovered dynamic shifts in specific cellular subpopulations following TAC surgery. Among the different immune cells in the heart, macrophages were the most abundant cell fraction indicating macrophages constituted the predominant immune subset in pathological cardiac hypertrophy (Figure 1K). Together, these data reveal a dynamic shift in cardiac cellular composition following pressure overload, with macrophages emerging as the predominant immune subset in the hypertrophic heart.

**Figure 1.**
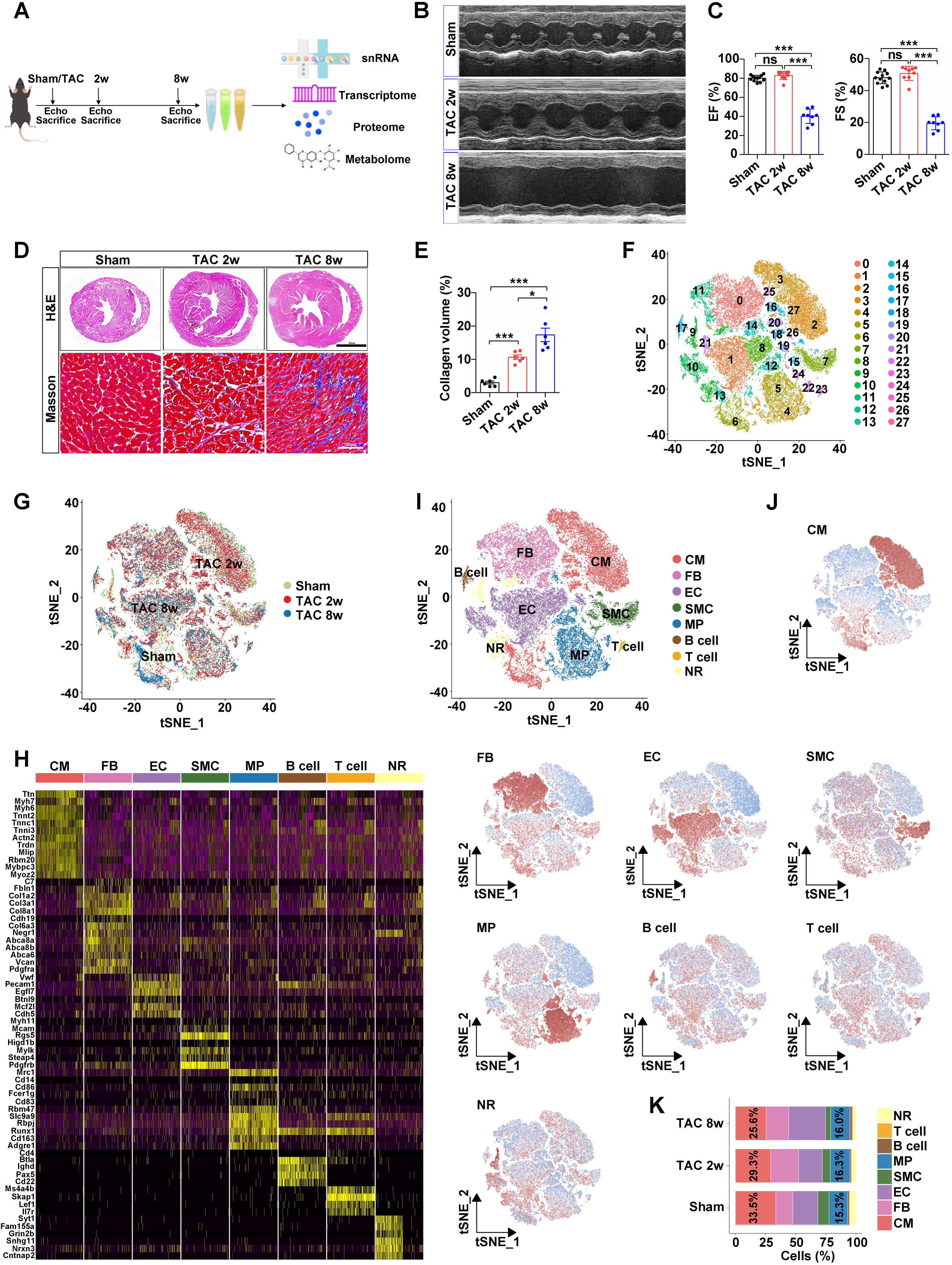
Single-nucleus analysis of pathological cardiac hypertrophy. **A,** Schematic overview of the study design. **B,** Representative M-mode echocardiographic images from Sham, 2 weeks post-TAC (TAC 2w), and 8 weeks post-TAC (TAC 8w) mice. **C**, Left ventricular ejection fraction (EF) and fractional shortening (FS) in mice after pressure overload (Sham, n=12; TAC 2w, n=9; TAC 8w, n=8). ***P<0.001; ns, not significant. D, Hematoxylin-eosin (H&E) staining (scale bar=2 mm) and Masson’s trichrome staining (scale bar=100 μm) were performed to determine the hypertrophic growth and fibrosis of the hearts from Sham, TAC 2w and TAC 8w mice. E, Quantification of cardiac fibrosis of the hearts from Sham, TAC 2w and TAC 8w mice (n=6). *P<0.05; ***P<0.001. F, t-SNE plot representing the 28 clusters across 53,957 cells from different stages of cardiac hypertrophy. G, t-SNE plot depicting 53,957 cells isolated from mouse hearts at different stages of cardiac hypertrophy. Each point represents a single cell, colored by cluster identity. **H,** Heat map showing the expression of marker genes in the indicated cell types. **I,** t-SNE plot showing the 8 annotated cell clusters. CM, cardiac muscle cells; FB, fibroblasts; EC, endothelial cells; SMC, smooth muscle cells; MP, macrophages; B, B cells; T, T cells; NR, neurogenic cells. **J,** t-SNE plot showing the distribution of 8 cell clusters identified in panel I. **K,** Bar chart showing the percentage distribution of each cell cluster in snRNA-seq datasets across different stages of pathological cardiac hypertrophy.

Myocardial metabolic remodeling is a core driving force in the progression from compensated cardiac hypertrophy to heart failure^19,20^. Compared to other cell types, cardiomyocytes exhibited the highest expression of glycolysis, tricarboxylic acid (TCA) cycle, and fatty acid oxidation (FAO) pathway genes (Figure S2D). To investigate the metabolic changes during pathological cardiac hypertrophy, the scores for the critical metabolic pathways in cardiomyocytes were calculated using single-sample Gene Set Enrichment Analysis (ssGSEA). This showed that glycolytic activity progressively increased while TCA cycle and FAO activity decreased post-TAC surgery (Figure S3A). Integrating transcriptome and proteome analysis confirmed the upregulation of the glycolytic regulators phosphofructokinase (Pfkp) and hexokinase 2 (Hk2), contrasting with the downregulation of bisphosphoglycerate mutase (Bpgm), aldehyde dehydrogenase 2 (Aldh2), and fructose-bisphosphatase 2 (Fbp2) (Figure S3B, C). Hk2 and Bpgm, in particular, serve as rate-limiting enzymes involved in regulating glucose metabolism, with HK2 controlling the first committed step of glycolysis by phosphorylating glucose, and BPGM modulating the 2,3-bisphosphoglycerate (2,3-BPG) shunt to balance glycolytic flux and redox homeostasis in mammalian cells^21–23^. The concomitant suppression of TCA metabolic genes, including malate dehydrogenase 1 (MDH1), dihydrolipoamide dehydrogenase (Dld), succinate dehydrogenase complex subunit B (Sdhb), succinate dehydrogenase complex subunit A (Sdha), and pyruvate dehydrogenase E1 subunit α1 (Pdha1), as well as FAO regulators such as medium-chain acyl-CoA dehydrogenase (Acadm), acetyl-CoA acyltransferase 2 (Acaa2), and 2,4-dienoyl-CoA reductase 1 (Decr1) were observed (Figure S3B, C). Metabolomic profiling indicated that differentially expressed metabolites involved in glycolysis, including glucose (Glc), glyceraldehyde-3-phosphate (GAP), fructose-1,6-bisphosphatase (FBP), and lactate were significantly up-regulated, indicating glycolysis was elevated to meet the enhanced energy demands during pathological cardiac hypertrophy (Figure S3D). In addition, TCA cycle–related metabolites such as malate and succinate peaked in concentration at 2 weeks post-TAC, followed by a decline at 8 weeks, while levels of 2-hydroxyglutarate (2-HG) and citrate decreased at 2 weeks but raised at 8 weeks after TAC surgery, consistent with stable and then declining cardiac function, implying the compensatory response of TCA cycle in pathological cardiac hypertrophy (Figure S3E). These results indicate that dynamic metabolic remodeling accompanies the progression of pathological cardiac hypertrophy.

### Cardiomyocyte subpopulations undergo dynamic remodeling during cardiac hypertrophy progression

Having established global immune and metabolic remodeling in hypertrophic hearts, we next focused on cardiomyocyte heterogeneity to identify subpopulations with distinct functional roles during disease progression. To delineate cardiomyocyte subpopulation dynamics in pathological cardiac hypertrophy, cardiomyocytes were divided into 14 transcriptionally distinct subtypes based on snRNA-seq data of Sham, TAC 2w and TAC 8w mice (Figure 2A, B). Pearson correlation-based functional clustering consolidated these into four cardinal functional clusters (FC) (CM FC1-4) with characteristic transcriptional programs (Figure 2C). CM FC1 highly expressed marker genes such as Cdh2 and participated in heart and muscle system processes, closely related to cardiac function (Figure 2D, E). CM FC2 was involved in the regulation of GTPase activity and highly expressed Ebf1, the main regulator of B cell development. Additionally, CM FC3 participated in ATP synthesis and cellular respiration and showed high expression of myoglobin. Interestingly, CM FC4 highly expressed typical fibroblast markers such as Col3a1 and Col1a1, which was functionally related to the regulation of extracellular structure organization. To identify the specific cardiomyocyte populations that may play a crucial role in pathological cardiac hypertrophy, we evaluated the proportions of cardiomyocyte subtypes and analyzed the dynamic changes in the distribution ratios of each subtype at different stages. CM FC1 accounted for the largest proportion within the cardiomyocyte population (Figure 2F, S4A). The cardiomyocyte subtypes exhibited dynamic changes over time: CM FC1 and FC2 were enriched in the early stages of cardiac hypertrophy, whereas CM FC3 and FC4 were mainly enriched in the late stages of cardiac hypertrophy (Figure 2G). To further understand the dynamic changes, pseudotime developmental trajectory analysis was carried out with CM subtypes (Figure 2H). CM FC1 was positioned at the forefront of the other subtypes, while CM FC2 was located relatively centrally, and CM FC3 and FC4 were mainly distributed at the end. In relative terms, CM FC1 had a higher differentiation potential, while CM FC3 and FC4 had lower differentiation potentials. Immunofluorescence co-staining of CDH2 and TNNT2 was performed to identify the CM FC1 subpopulation within the heart tissues of mice (Figure 2I). Therefore, these results elucidate the cardiomyocyte subpopulations and their potential roles during pathological cardiac hypertrophy.

**Figure 2.**
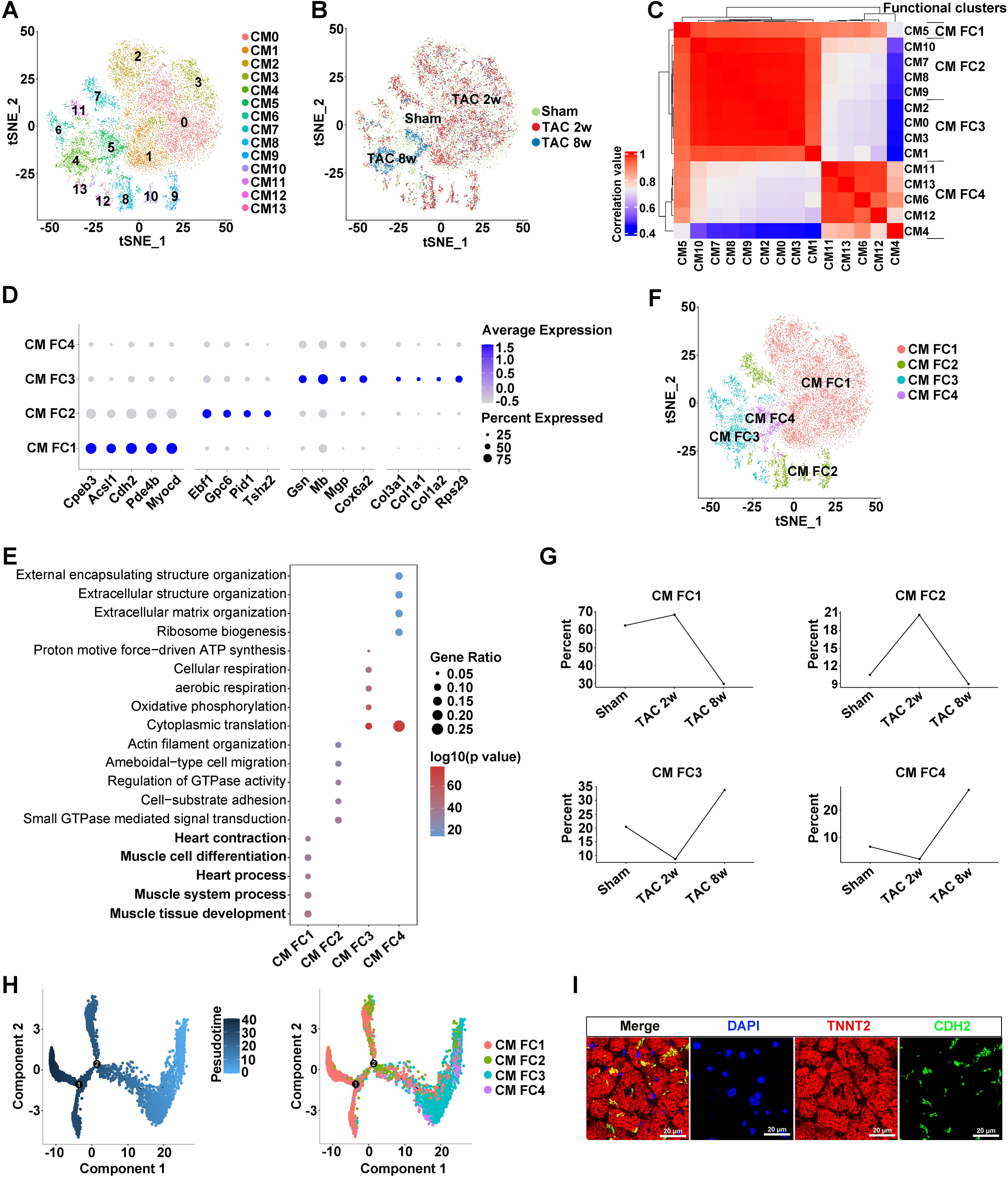
Single-nucleus RNA sequencing reveals cardiomyocyte subtypes at different stages of cardiac hypertrophy. **A-B,** t-SNEs plot showing 16,391 cardiomyocytes (CM) isolated from mouse hearts at different stages of cardiac hypertrophy. Cells were marked by cluster number (A) or subcluster number (B). C, 14 identified CM clusters were categorized into functional clusters (FCs) utilizing Spearman’s rank correlation coefficient analysis. FC1=CM5, FC2=CM7+CM8+CM9+CM10, FC3=CM0+CM1+CM2+CM3, FC4=CM4+CM6+CM11+CM12+CM13. D, Bubble chart showing the expression of marker genes for each CM FC. E, Gene ontology analysis of specifically expressed genes in each CM FC. F, t-SNE projection of cardiomyocyte subtypes after unsupervised clustering. G, Changes in the proportions of CM subtypes at different stages of cardiac hypertrophy, normalized to total number of CMs at each time point. **H,** Pseudo-time trajectories of the CM subtypes inferred by Monocle trajectory analysis. **I,** Immunofluorescence staining for TNNT2 (red) and CDH2 (green) on heart sections 2 weeks after TAC surgery. Scale bar=20 μm.

### Distinct macrophage subpopulations shift dynamically and reveal a CD163⁺/Dab2⁺ subset linked to disease progression

Given our earlier finding that macrophages are the predominant immune population in the hypertrophic heart, we next sought to analyze the heterogeneity and dynamic changes of macrophages during disease progression. Twelve macrophage subpopulations were identified, including MP0-11 by single-nucleus transcriptional profiling of hearts from Sham, TAC 2w and TAC 8w mice (Figure 3A, B), and then consolidated into five functionally distinct clusters (MP FC1-5) based on their similar transcriptional profiles through Pearson correlation analysis (Figure 3C). MP FC1, highly expressed Tmtc2 and Adgrl4, was primarily related to insulin signaling and chemokine signaling pathway (Figure 3D, E). MP FC2 significantly expressed M2-like markers CD163, F13a, and the tumor suppressor Dab2^24–26^ (Figure 3D). As a novel regulators of macrophage phenotypic polarization, Dab2 expression was upregulated in M2 macrophages and suppressed in M1 macrophages isolated from both mice and humans, and genetic deletion of Dab2 predisposed macrophages in mice to adopt a proinflammatory M1 phenotype^27^. Functional analysis revealed that MP FC2 was enriched mainly in immunity-related processes, including Kit receptor signaling, IL-3 signaling and IL-5 signaling pathways (Figure 3E). MP FC3 significantly expressed Ebf1 and Mast4 and were significantly correlated with focal adhesion and endochondral ossification. MP FC4 was mainly involved in biological processes such as electron transport chain and oxidative phosphorylation, while MP FC5 participated in amino acid metabolism and fatty acid beta oxidation. Among these macrophages, MP FC1 and MP FC2 were the most abundant subtypes in the hearts during pathological cardiac hypertrophy (Figure 3F, S4B). Moreover, the macrophage subpopulations exhibited distinct distribution patterns in pathological cardiac hypertrophic hearts. In response to disease, the relative abundance of MP FC1 was reduced in TAC-operated mice in comparison with Sham, whereas MP FC5 was progressively increased (Figure 3G). The relative abundances of MP FC2 and FC4 were declined in 2 weeks after TAC surgery but were elevated 8 weeks post-TAC. Conversely, the abundance of MP FC3 peaked at 2 weeks after TAC and subsequently decreased at 8 weeks. The pseudotime developmental trajectory analysis revealed the dynamic trend in macrophages (Figure 3H). MP FC2 was mainly positioned at the forefront of other subtypes; MP FC1, FC4 and FC5 were located relatively centrally; while MP FC3 was mainly distributed at the end, suggesting the potential involvement of these subpopulations in cardiac hypertrophy. Relatively, MP FC2 had higher differentiation potential, while MP FC3 and FC4 had lower differentiation potential. noting its M2-like marker expression and possible regulatory function. Notably, these distinctions may correlate with the M2-like marker expression profile of MP FC2 and suggests its potential regulatory role in immune modulation. Furthermore, immunofluorescence co-staining of CD163 and Dab2 was performed to identify the MP FC2 subtype within the heart tissue of mice (Figure 3I). Altogether, these findings provide evidence that the CD163^+^/Dab2^+^ macrophages subtype may play an active role in the progression of pathological cardiac hypertrophy.

**Figure 3.**
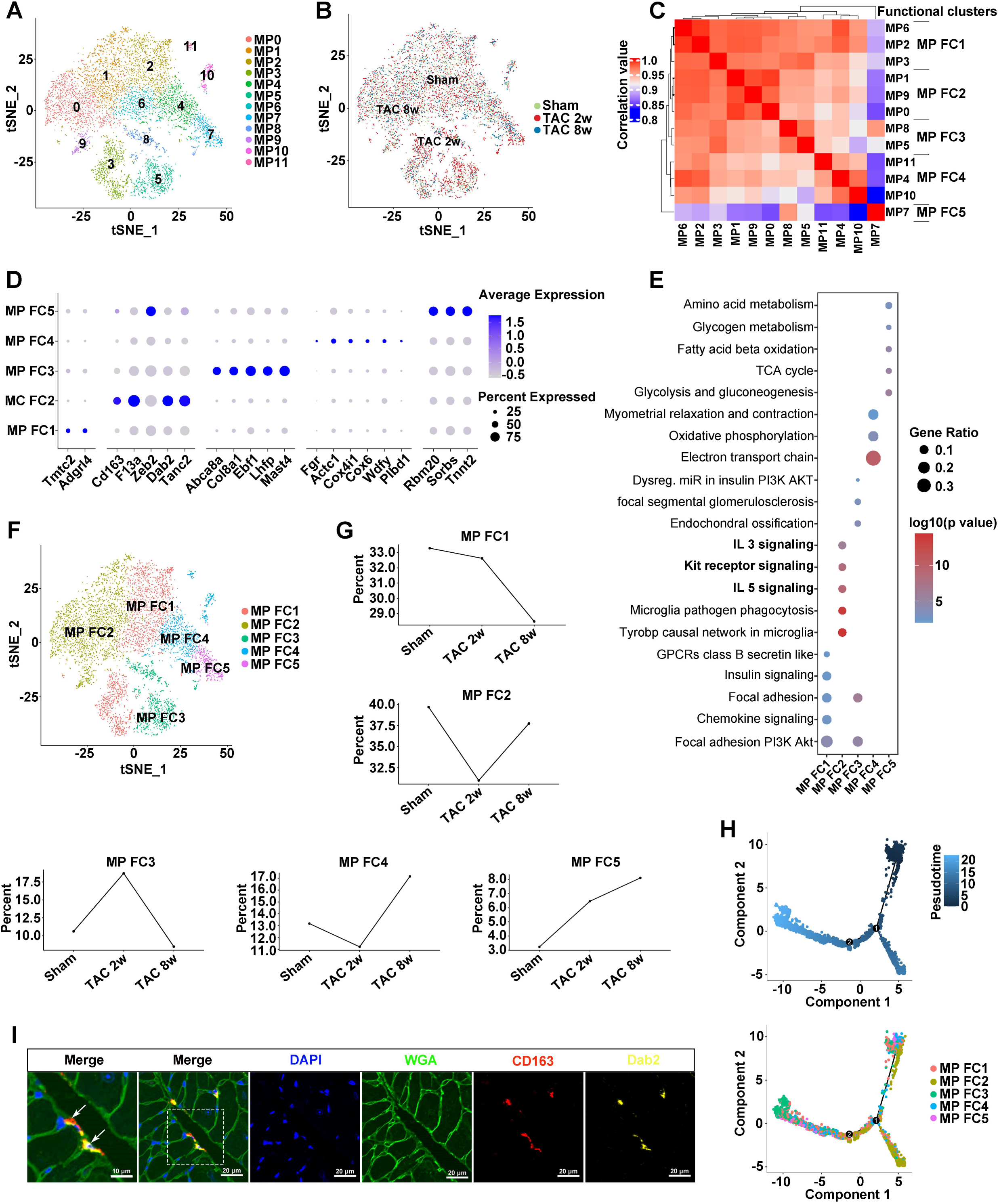
Characterization of macrophage subtypes at different stages of pathological cardiac hypertrophy. **A-B,** t-SNE plots showing 8,553 macrophages (MPs) isolated from mouse hearts at different stages of cardiac hypertrophy. Cells were marked by cluster number (A) or subcluster number (B). C, 12 identified MP clusters were categorized into functional clusters (FCs) utilizing Spearman’s rank correlation coefficient analysis. FC1=MP2+MP3+MP6, FC2=MP0+MP1+MP9, FC3=MP5+MP8, FC4=MP4+MP10+MP11, FC5=MP7. D, Bubble chart showing the expression of marker genes for each MP FC. E, WikiPathways analysis of specifically expressed genes in each MP FC. F, t-SNE projection of MP subtypes after unsupervised clustering. G, Changes in proportions of MP subtypes at different stages of cardiac hypertrophy, normalized to total number of MPs at each time point. **H,** Pseudo-time trajectories of the MP subtypes inferred by Monocle trajectory analysis. **I,** Wheat germ agglutinin (WGA) staining (green) and immunofluorescence staining against CD163 (red) and Dab2 (yellow) on heart sections 2 weeks after TAC surgery. Scale bar=20 μm.

### CD163^+^/Dab2^+^ macrophages communicate with cardiomyocytes via the Nrg2/ErbB4 pathway

Given the observed shifts in macrophage populations and their proximity to cardiomyocytes, we next investigated whether specific macrophage subtypes engage in direct intercellular communication with cardiomyocytes. To evaluate differences in the molecular interactions between cells during pathological cardiac hypertrophy, we used CellPhoneDB^28^ to construct the intercellular communication networks based on known ligand‒receptor pairs and their accessory factors (Figure 4A). The detection of cognate receptors for broadcast ligands within both cardiomyocytes and cardiac non-myocytes revealed that macrophages form one of the most trophic cell populations, showing dense connections with cardiomyocytes (Figure 4A). Expression patterns of the ligand-receptor pairs between macrophages and other cell types revealed a dense intercellular communication network. Specifically, macrophages closely communicated with cardiomyocytes by the ligand-receptor pairs Nrg2/ErbB4 in the hearts of Sham mice (Figure 4B), whereas macrophages interacted with cardiomyocytes via other ligand-receptor pairs in the hearts of mice at 2 weeks and 8 weeks post-TAC surgery, including FGF1/FGFR2 and TF/TFRC, respectively (Figure S5A, B). This shift in interactions may be attributed to the decreased expression of the Nrg2/ErbB4 ligand-receptor pair. Interestingly, the MP FC2 subtype (CD163^+^/Dab2^+^) also tightly communicated with cardiomyocytes through the ligand-receptor pairs Nrg2/ErbB4 in the hearts of Sham mice (Figure 4C). Furthermore, the ligand Nrg2 and receptor ErbB4 were expressed at the highest levels in the MP FC2 macrophage subpopulation and CM FC1 cardiomyocyte subpopulation, respectively (Figure S5C, D). To investigate the dynamic expression of Nrg2 and ErbB4, we utilized single-nucleus transcriptomics to reveal that they were downregulated in macrophages and cardiomyocytes, respectively (Figure 4D). In addition, both bulk-RNA sequencing and RT-qPCR demonstrated the reduced expression of Nrg2 and ErbB4 in mouse hearts during pathological cardiac hypertrophy (Figure 4E, F). Likewise, scRNA-seq datasets from the Single Cell Portal under project ID SCP1303^29^, comprising 16 normal human heart samples, 11 dilated cardiomyopathy human heart (DCM) samples and 15 hypertrophic cardiomyopathy human heart (HCM) samples were analyzed, and the results suggested that the levels of Nrg2 and ErbB4 were diminished in DCM and HCM compared to Normal samples (Figure S5E, F). Immunofluorescence staining of mouse hearts after TAC confirmed the colocalization of TNNT2^+^/CDH2^+^ cardiomyocytes with ErbB4 and the colocalization of CD163^+^/Dab2^+^ macrophages with Nrg2 (Figure 4G, H). Additionally, we observed the colocalization of TNNT2 and ErbB4 in neonatal rat ventricular myocytes (NRVMs) and colocalization of CD163 and Nrg2 in bone marrow-derived macrophages (BMDMs) (Figure S5G, H). The intercellular communication of CD163^+^/Dab2^+^ macrophages and cardiomyocytes through the ligand-receptor pair Nrg2/ErbB4 was validated by multiplex immunofluorescence staining (Figure 4I). Further, co-immunoprecipitation confirmed the interaction between Nrg2 and ErbB4 in NRVMs (Figure 4J). Therefore, these findings suggest that CD163^+^/Dab2^+^ macrophages communicate with cardiomyocytes through the Nrg2/ErbB4 pathway in the heart.

**Figure 4.**
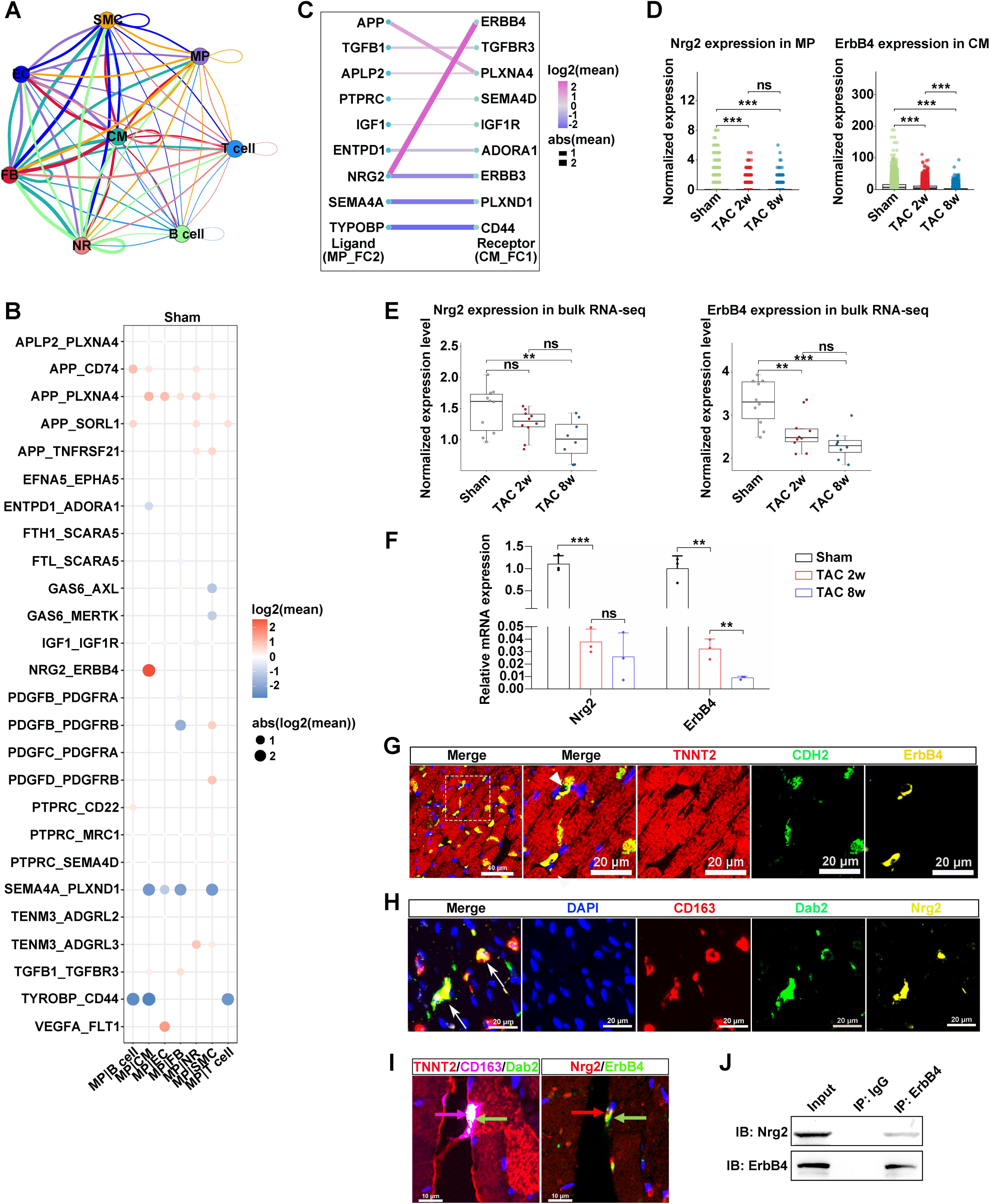
CD163^+^/Dab2^+^ macrophages communicate with cardiomyocytes via the Nrg2/ErbB4 pathway. **A,** Schematic illustration of intercellular communication among cardiac cell types, with line colors representing ligands produced by the corresponding cell populations. Lines connected to those cell groups that expressed compatible receptors, with thickness indicating the relative abundance of ligands compared to receptor presence. Looping lines highlighted autocrine signaling. CM, cardiac muscle cells; FB, fibroblasts; EC, endothelial cells; SMC, smooth muscle cells; MP, macrophages; NR, neuronal cells. **B,** Putative ligand-receptor pairs linking macrophages (MP) to other cardiac cell types in Sham hearts. **C,** Putative ligand-receptor pairs between MP FC2 and CM FC1 in hearts from Sham mice. **D,** Single-nucleus RNA sequencing data for Nrg2 expression in macrophages and ErbB4 expression in cardiomyocytes at different stages of cardiac hypertrophy. \*\*\**P*<0.001; ns, not significant. **E,** Bulk RNA sequencing analysis of Nrg2 and ErbB4 expression at different stages of cardiac hypertrophy (Sham, n=10; TAC 2w, n=10; TAC 8w, n=8). \*\**P*<0.01; \*\*\**P*<0.001; ns, not significant. **F,** Relative mRNA expression levels of Nrg2 and ErbB4 in mouse hearts at different stages of cardiac hypertrophy, as analyzed by RT-qPCR. (n=3). \*\**P*<0.01; \*\*\**P*<0.001; ns, not significant. **G,** Multicolor immunofluorescence staining for TNNT2 (red), CDH2 (green), and ErbB4 (yellow) in heart sections 2 weeks after TAC surgery. Nuclei were counterstained with DAPI (blue). White arrowhead indicates the colocalization of TNNT2^+^/CDH2^+^ cardiomyocytes with the receptor ErbB4. Scale bar=20 μm. **H,** Multicolor immunofluorescence staining for CD163 (red), Dab2 (green), and Nrg2 (yellow) in heart sections 2 weeks after TAC surgery. Nuclei were counterstained with DAPI (blue). White arrow indicates the colocalization of CD163^+^/Dab2^+^ macrophages with the ligand Nrg2. Scale bar=20 μm. **I,** Multicolor immunofluorescence staining for TNNT2 (red), CD163 (pink), Dab2 (green), or Nrg2 (red) and ErbB4 (green) in heart sections 2 weeks after TAC surgery. Nuclei were counterstained with DAPI (blue). **Left,** pink arrow indicates CD163^+^ field, and green arrow indicates Dab2^+^ field. **Right,** red arrow indicates Nrg2^+^ field, and green arrow indicates ErbB4^+^ field. Scale bar=10 μm. **J,** Co-immunoprecipitation (Co-IP) analysis of the interactions between Nrg2 and ErbB4 in neonatal rat ventricular myocytes (NRVMs).

### CD163^+^/Dab2^+^ macrophages assuage cardiac hypertrophy via the Nrg2/ErbB4 pathway in vitro

To determine whether CD163^+^/Dab2^+^ macrophages can modulate pathological cardiac hypertrophy in vitro, we sorted CD163^+^/Dab2^+^ macrophages and confirmed their positive correlation with Nrg2 expression by flow cytometry (Figure 5A). To evaluate differential Nrg2 secretion among macrophage subtypes, conditioned medium from FACS-sorted CD163^-^, CD163^+^/Dab2^-^, and CD163⁺/Dab2⁺ macrophages were analyzed by ELISA, which revealed elevated levels of Nrg2 in CD163⁺/Dab2⁺ macrophages compared to CD163^-^ and CD163^+^/Dab2^-^ subsets (Figure 5B). Conditioned medium from CD163^+^/Dab2^+^ macrophages mitigated cardiac hypertrophy induced by Ang II and significantly reduced the expression of hypertrophy-related genes, such as Anp and Myh7 (Figure 5C-E). On the other hand, when the expression of Nrg2 was knocked down in CD163^+^/Dab2^+^ macrophages by shRNA (Figure S6A, B), we found that conditioned medium from CD163^+^/Dab2^+^ macrophages with Nrg2 knockdown significantly promoted cardiac hypertrophy and upregulated the expression of Anp and Myh7 (Figure 5C-E). To further determine whether CD163^+^/Dab2^+^ macrophages regulate cardiac hypertrophy via the Nrg2/ErbB4 pathway, NRVMs were treated with recombinant Nrg2 and the ErbB4 inhibitor AG1478. Recombinant Nrg2 promoted the level of pErbB4 in NRVMs, proving the pathway was activated, while AG1478 inhibited its phosphorylation (Figure S6C). In particular, Nrg2 treatment prominently diminished the mRNA and protein levels of Anp and Myh7, and the enlarged cardiomyocyte size caused by Ang II, whereas the addition of AG1478 rescued the inhibitory effect of Nrg2 on cardiac hypertrophy in vitro (Figure 5F-I). Moreover, the administration of Nrg2 reduced the cardiomyocyte apoptosis caused by Ang II (Figure S6D, E). Collectively, these results indicate that CD163^+^/Dab2^+^ macrophages alleviate cardiac hypertrophy via the Nrg2/ErbB4 pathway in vitro.

**Figure 5.**
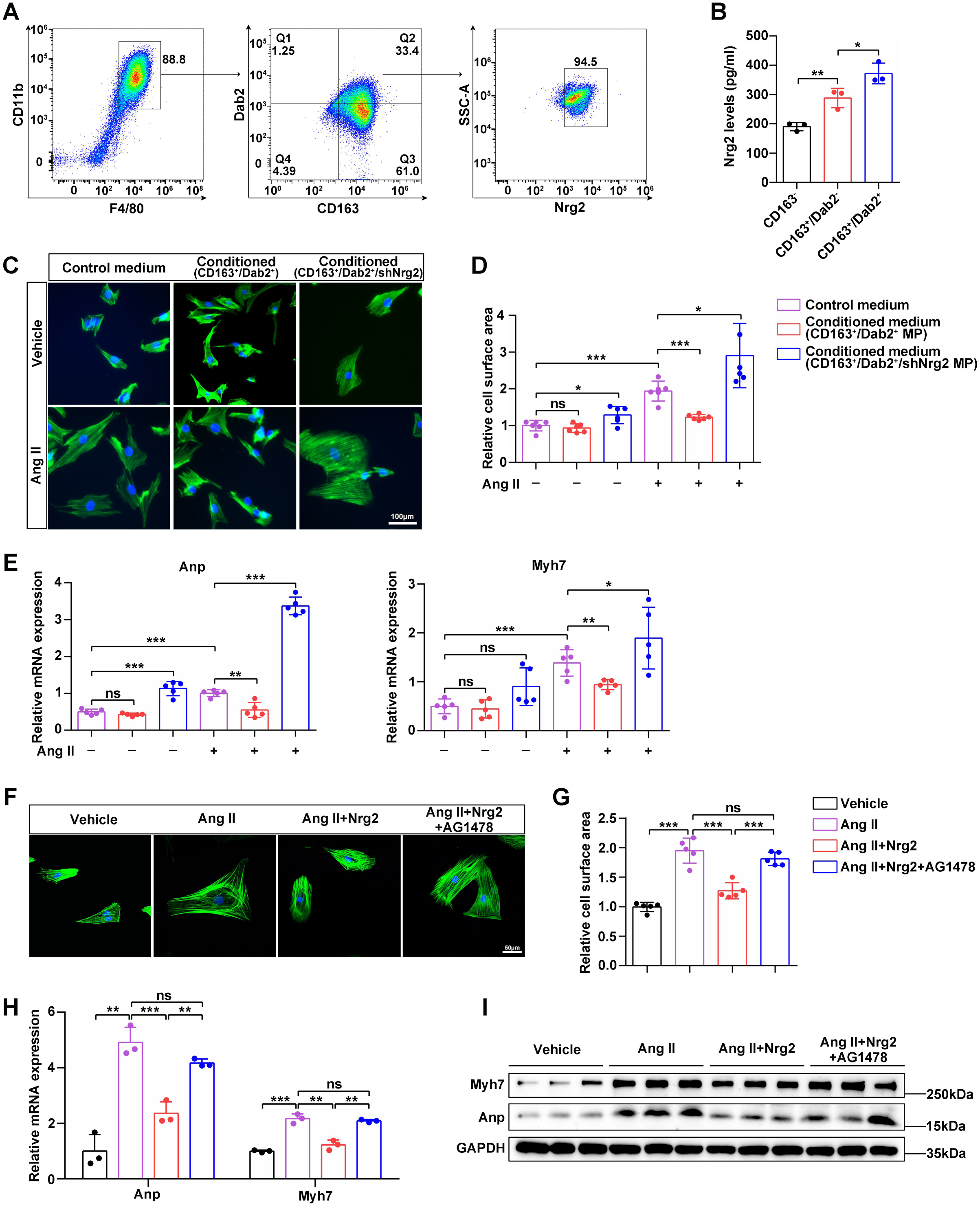
CD163^+^/Dab2^+^ macrophages mitigate cardiac hypertrophy via the Nrg2/ErbB4 pathway in vitro. **A,** Representative flow plots of F4/80, CD11b, CD163, Dab2 and Nrg2 expression in polarized BMDMs. **B,** ELISA analysis of Nrg2 levels in CD163^-^, CD163^+^/Dab2^-^, and CD163^+^/Dab2^+^ macrophages (n=3). \**P*<0.05; \*\**P*<0.01. **C,** Representative immunofluorescence images of phalloidin staining in NRVMs cultured with different medium and treated with vehicle or Ang II for 48 hours. Scale bar=100 μm. **D,** Relative cell surface area of NRVMs cultured with different medium following vehicle or Ang II treatment (n=6). \**P*<0.05; \*\*\**P*<0.001; ns, not significant. **E,** RT-qPCR analysis of the mRNA levels of Anp and Myh7 in NRVMs cultured with different medium and treated with vehicle or Ang II for 48 hours (n=5). \**P*<0.05; \*\**P*<0.01; \*\*\**P*<0.001; ns, not significant. **F,** Representative immunofluorescence images of phalloidin staining in NRVMs treated with recombinant Nrg2 and inhibitor AG1478 in response to vehicle or Ang II treatment for 48 hours, respectively. Scale bar=100 μm. **G,** Relative cell surface area of NRVMs treated with recombinant Nrg2 and/or inhibitor AG1478 in response to vehicle or Ang II treatment for 48 hours, respectively (n=5). \*\*\**P*<0.001; ns, not significant. **H,** RT-qPCR analysis of the mRNA levels of Anp, Bnp and Myh7 in NRVMs treated with recombinant Nrg2 and and inhibitor AG1478 in response to vehicle or Ang II treatment for 48 hours, respectively (n=3). \**P*<0.05; \*\**P*<0.01; \*\*\**P*<0.001. **I,** Protein expression of Anp and Myh7 in NRVMs treated with recombinant Nrg2 and inhibitor AG1478 in response to vehicle or Ang II treatment for 48 hours, respectively

### Nrg2/ErbB4 signaling attenuates cardiac hypertrophy in vivo

To further explore the role of Nrg2/ErbB4 pathway in cardiac hypertrophy, recombinant Nrg2 and the inhibitor AG1478 were injected into mice 1 week after TAC surgery (Figure 6A). The heart weight/body weight ratio of mice in TAC+Nrg2 group was lower than that of mice in TAC group and TAC+Nrg2+AG1478 group 8 weeks after TAC surgery, whereas there was no difference between mice in TAC group and mice in TAC+Nrg2+AG1478 group (Figure 6B, C). TAC surgery resulted in the reduced cardiac output (CO), ejection fraction (EF) and fractional shortening (FS), but these were increased in Nrg2-treated mice (Figure 6D, E), indicating that Nrg2 had a protective effect on cardiac function. However, the simultaneous administration of Nrg2 and AG1478 did not increase cardiac output, ejection fraction or fractional shortening after TAC surgery (Figure 6D, E). Additionally, morphological analyses showed the heart size and cross-sectional area (CSA) of mice in TAC+Nrg2 group were significantly less than those of mice in TAC group and TAC+Nrg2+AG1478 group, but the difference between mice in TAC+Nrg2+AG1478 group and TAC group was not significant (Figure 6F, G). Moreover, TAC surgery significantly upregulated the mRNA expression of cardiac hypertrophy markers and fibrosis markers, including Anp, Bnp, Myh7, Col1a1, Col3a1 and Ctgf, whereas their expression levels were dramatically declined in Nrg2-treated mice (Figure 6H, I). Compared to mice in TAC+Nrg2 group, the mRNA expression of cardiac hypertrophy markers and fibrosis markers were upregulated in mice with simultaneous administration of Nrg2 and AG1478 (Figure 6H, I). Consistently, administration of recombinant Nrg2 substantially attenuated the fibrosis resulted from TAC surgery, while the fibrosis of mice in TAC+Nrg2+AG1478 group was remarkably higher than mice in TAC+Nrg2 group (Figure 6J, K). On the other hand, the rate of apoptotic cells was notably reduced in Nrg2-treated TAC mice compared to TAC only mice (Figure S6F, G). Accordingly, these data establish Nrg2/ErbB4 activation as a viable strategy for mitigating pathological cardiac hypertrophy.

**Figure 6.**
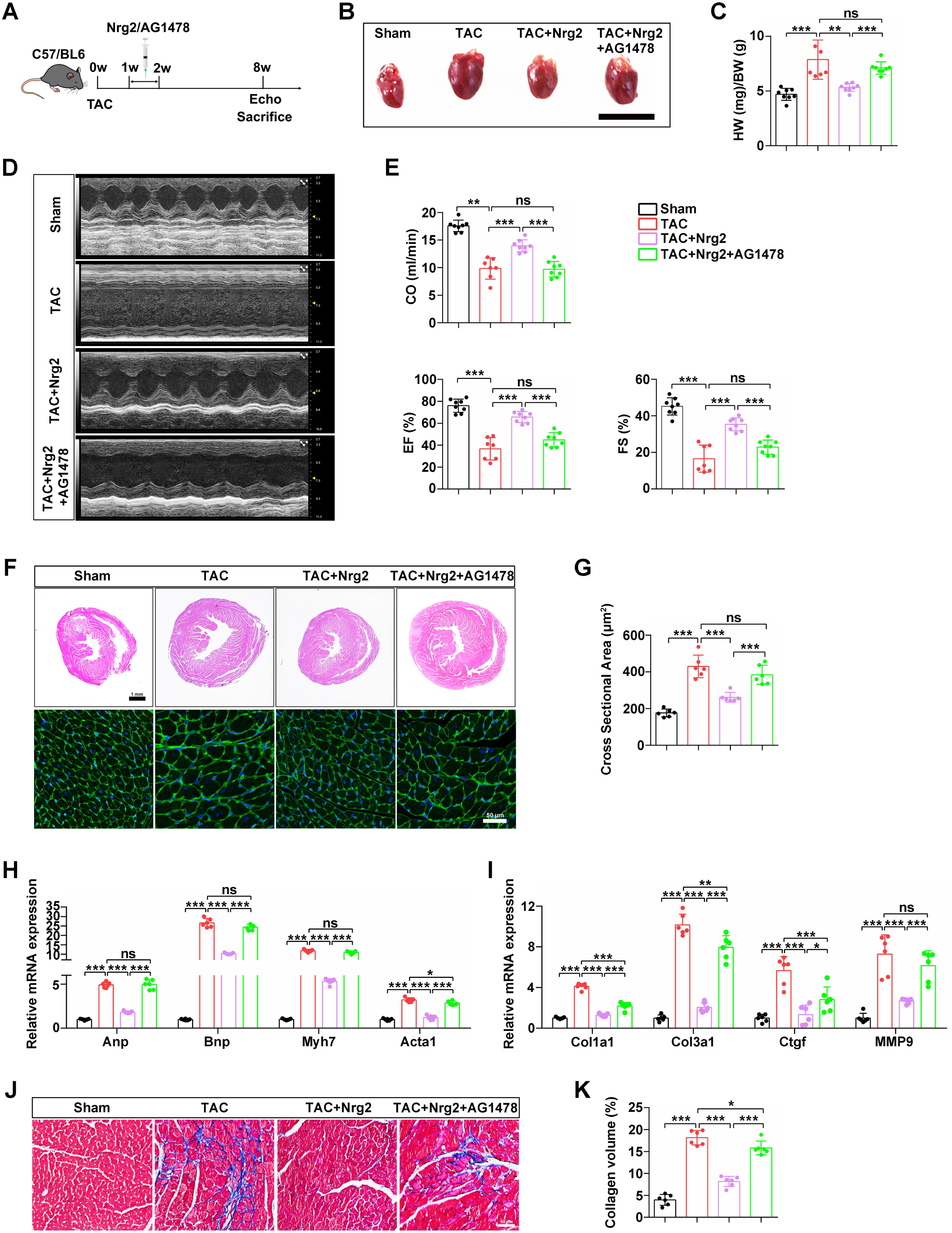
Activation of Nrg2/ErbB4 pathway alleviates cardiac hypertrophy in vivo. **A,** Schematic diagram illustrating the experimental design involving the injection of recombinant Nrg2 and inhibitor AG1478 into a mouse model of cardiac hypertrophy. Wildtype mice at 6 weeks of age were subjected to transverse aortic constriction (TAC) or Sham surgery. After 1 week, recombinant Nrg2 or Nrg2 and inhibitor AG1478 were continuously administered into the mice for 7 days. These mice groups were denoted as Sham, TAC, TAC+Nrg2, and TAC+Nrg2+AG1478. **B,** Representative images of hearts at 8 weeks post-surgery from mice subjected to Sham or TAC surgery and injected with recombinant Nrg2 or Nrg2 and inhibitor AG1478. **C,** Heart weight/body weight (HW/BW) ratios at 8 weeks post-surgery for mice subjected to Sham or TAC surgery and injected with recombinant Nrg2 or Nrg2 and inhibitor AG1478 (Sham, n=8; TAC, n=7; TAC+Nrg2, n=8; TAC+Nrg2+AG1478, n=8). \*\**P*<0.01; \*\*\**P*<0.001; ns, not significant. **D and E,** M-mode echocardiography images (D) and quantified parameters (E) of the left ventricular (LV) chamber at 8 weeks post-surgery in mice subjected to Sham or TAC surgery and injected with recombinant Nrg2 or Nrg2 and inhibitor AG1478 (Sham, n=8; TAC, n=7; TAC+Nrg2, n=8; TAC+Nrg2+AG1478, n=8). \*\**P*<0.01; \*\*\**P*<0.001; ns, not significant. EF, ejection fraction; FS, fractional shortening; CO, cardiac output. **F,** Hematoxylin-eosin (H&E) (scale bar=1 mm) and Wheat germ agglutinin (WGA) staining (scale bar=50 μm) showing morphology and cell boundaries of heart sections 8 weeks after TAC with indicated treatment regimens. **G,** Quantification of the cardiomyocyte cross-sectional areas of the indicated groups (n=6 mice per group). \*\*\**P*<0.001; ns, not significant. **H,** The mRNA levels of hypertrophic marker genes Anp, Bnp, Myh7 and Acta1 in hearts of the indicated groups (n=6 mice per group). \**P*<0.05; \*\*\**P*<0.001; ns, not significant. **I,** The mRNA levels of fibrosis marker genes Col1a1, Col3a1, Ctgf and MMP9 in hearts of the indicated groups (n=6 mice per group). \**P*<0.05; \*\**P*<0.01; \*\*\**P*<0.001; ns, not significant. **J,** Masson’s trichrome staining showing fibrosis in heart sections 8 weeks after TAC with indicated treatment regimens. Scale bar=50 μm. **K,** Quantification of cardiac fibrosis in tissue shown in panel J (n=6 mice per group). \**P*<0.05; \*\*\**P*<0.001.

### Recombinant Nrg2 restores mitochondrial bioenergetics in hypertrophic hearts

To explore the mechanisms underlying the protective effects of Nrg2/ErbB4 signaling, we performed transcriptomic analysis of heart tissue from Sham, TAC and Nrg2-treated TAC mice (Figure 7A). Compared with Sham mice, 1,449 genes were upregulated and 1,367 genes were downregulated in the hearts of TAC only mice, and 945 genes were upregulated and 887 genes were downregulated in the hearts of Nrg2-treated TAC mice based on fold changes (>1.0) and P values (<0.05) (Figure S7A). 1,229 upregulated differentially expressed genes and 1,422 downregulated differentially expressed genes were identified in Nrg2-treated TAC mice compared to TAC-only mice (Figure 7B). As anticipated, the expression of cardiac hypertrophy and fibrosis genes, including Nppa, Nppb, Acta1, Col1a2, Col3a1 and Postn, were decreased in Nrg2-treated TAC mice compared to TAC mice (Figure 7C). Functional annotation showed that upregulated genes in Nrg2-treated TAC mice were mainly linked to muscle contraction and cell differentiation, while in TAC-only mice, upregulated genes were associated with extracellular matrix organization and muscle cell differentiation (Figure S7B). In particular, these differentially expressed genes between Nrg2-treated TAC mice and TAC only mice were significantly enriched for mitochondrial metabolic pathways including mitochondrial ATP synthesis, oxidative phosphorylation and cellular respiration (Figure 7D). Likewise, Gene Set Enrichment Analysis (GSEA) revealed that oxidative phosphorylation activity levels were raised in Nrg2-treated TAC mice compared with TAC only mice (Figure 7E, S7C). Mitochondrial oxidative phosphorylation is a major energy source for the normal heart, and reduced oxidative phosphorylation is closely related to mitochondrial dysfunction^30,31^. The expression levels of mitochondrial respiratory electron transport chain (ETC) genes were analyzed by RNA-sequencing which were broadly upregulated in Nrg2-administered mice after TAC surgery (Figure 7F). Moreover, the mRNA expression of ETC complex subunits in hearts was verified by RT-qPCR and was found to be lower in the hearts of TAC mice than those of Sham mice; however, the expression of these ETC genes was obviously enhanced in the hearts of Nrg2-treated TAC mice compared to TAC mice (Figure 7G). Taken together, our results reveal that the protective effect of Nrg2 in hypertrophic hearts is mediated by restoring mitochondrial oxidative metabolism.

**Figure 7.**
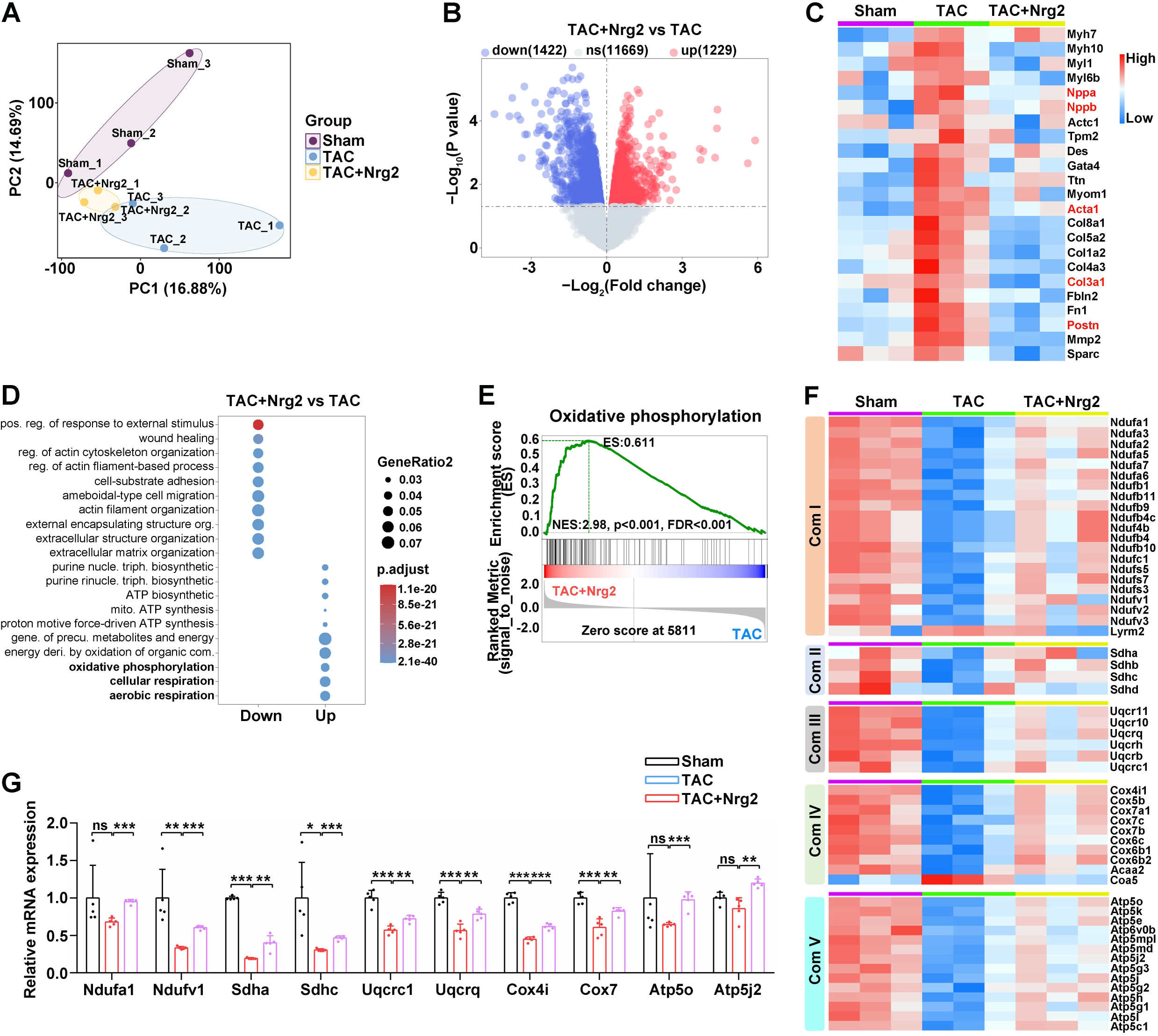
Administration of Nrg2 alleviates cardiac hypertrophy by improving mitochondrial bioenergetics. **A,** Principal component analysis (PCA) plot illustrating the transcriptional variance among hearts in the indicated groups. **B,** Volcano plot analysis showing the differential genes between TAC and TAC+Nrg2 group. **C,** Heatmap displaying the differentially expressed hypertrophic marker genes and fibrosis marker genes in hearts from Sham, TAC, and Nrg2+Nrg2 mice at 8 weeks post-surgery. **D and E,** GO enrichment analysis (D) and gene set enrichment analysis (E) of RNA-sequence data from Sham, TAC, and Nrg2+Nrg2 mice hearts at 8 weeks post-surgery. **F,** Heatmap showing the expression of mitochondrial respiratory electron transport chain (ETC, Complex I–V) genes. **G,** The mRNA expression of ETC complex genes in Sham, TAC, and TAC+Nrg2 group hearts 8 weeks after TAC surgery (n=5 for each group). \**P*<0.05; \*\**P*<0.01; \*\*\**P*<0.001; ns, not significant.

### CD163^+^/Dab2^+^ macrophages restore the mitochondrial dysfunction induced by Ang II via the Nrg2/ErbB4 pathway

To further examine the role of the Nrg2/ErbB4 pathway in cardiac mitochondrial function, we analyzed the ultrastructure of mitochondria in the hearts of mice using transmission electron microscopy, and found that TAC surgery increased mitochondrial size and caused cristae disorganization and mitochondrial damage (Figure 8A, B). Conversely, Nrg2 administration rescued the abnormal mitochondrial morphology and density that resulted from TAC surgery (Figure 8A, B). In addition, the ATP content in TAC mouse hearts was depressed compared to Sham mouse hearts, but it was higher in Nrg2-treated TAC mice than TAC mice, indicating Nrg2 improved mitochondrial bioenergetics in vivo (Figure 8C). Mitochondrial dysfunction and disruptions in homeostasis are critical factors in diminishing myocardial contractility and are closely linked to the development and progression of heart failure. Maintaining mitochondrial homeostasis safeguards cardiomyocytes from damage and preserves cardiac function. Dysregulation of mitochondrial fusion, fission, and mitophagy leads to cardiomyocyte dysfunction or death, thereby accelerating the onset and exacerbation of heart failure^12,32^. To investigate the effect of Nrg2 on mitochondrial function in vitro, NRVMs were treated with Ang II or Ang II and recombinant Nrg2 in vitro for 48 h, then assessed by Mitotracker Red. Nrg2 administration mitigated the Ang II-induced abnormalities in mitochondrial distribution and morphology, and decreased the frequency of mitochondrial fission in NRVMs (Figure 8D, E). Furthermore, extracellular flux analysis was performed to real-time monitor the mitochondrial respiratory capacity in vitro. Nrg2 administration restored the mitochondrial respiratory capacity after Ang II treatment, with observed increases in basal respiration, ATP production coupled respiration and maximal respiration (Figure 8F, G). Analogously, we evaluated the impact of CD163^+^/Dab2^+^ macrophages on cardiomyocyte mitochondrial function by extracellular flux analysis. The results revealed that conditioned medium from CD163^+^/Dab2^+^ macrophages improved the mitochondrial respiratory capacity after Ang II treatment, with observed increases in basal respiration, ATP production coupled respiration and maximal respiration (Figure 8H, I). Contrarily, the culture of cardiomyocytes in conditioned medium derived from Nrg2-knockdown CD163^+^/Dab2^+^ macrophages led to a significant reduction in mitochondrial respiratory capacity, with reduced basal respiration, ATP-linked respiration, and maximal respiration (Figure 8H, I). Moreover, Mitotracker Red staining implied that conditioned medium from CD163^+^/Dab2^+^ macrophages rescued the mitochondrial morphological abnormalities caused by Ang II, whereas conditioned medium from Nrg2-knockdown CD163^+^/Dab2^+^ macrophages did not have a protective effect on mitochondrial morphology in cardiomyocytes (Figure 8J, K). These data demonstrate that CD163^+^/Dab2^+^ macrophages rescued Ang II-induced mitochondrial dysfunction in cardiomyocytes via the Nrg2/ErbB4 pathway in vitro.

**Figure 8.**
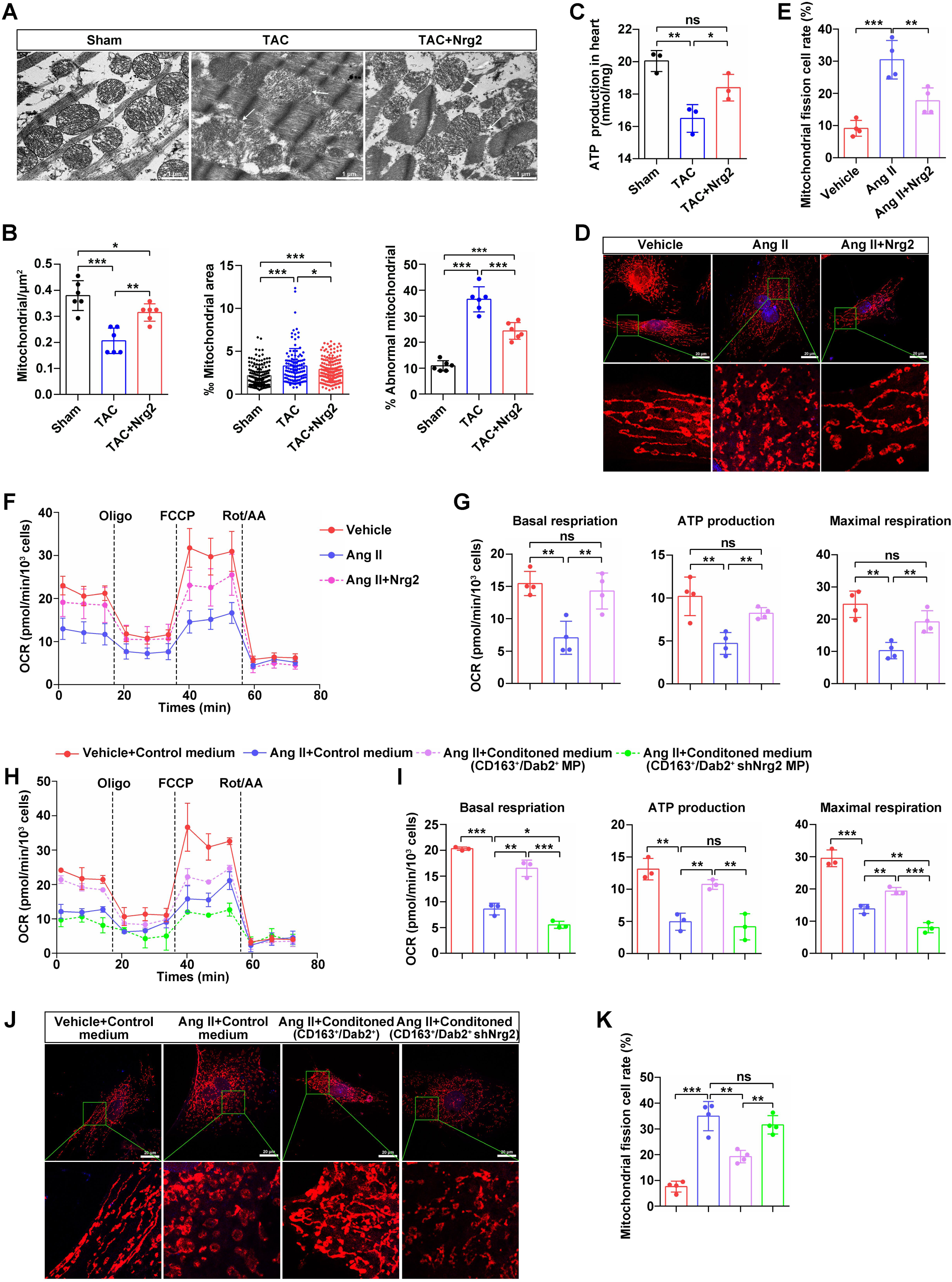
CD163^+^/Dab2^+^ macrophages restore cardiac mitochondrial function via the Nrg2/ErbB4 pathway. **A,** Representative transmission electron microscopy images of heart tissues from Sham, TAC and TAC+Nrg2 group mice at 8 weeks post-TAC surgery. Scale bar=1 μm. **B,** Quantification of mitochondria-related parameters in A. Mitochondria density per area (mitochondria/μm²) (n=6 for each group), ratio of mitochondrial area to image area (‰ mitochondrial area) (Sham, n=148; TAC, n=118; TAC+Nrg2, n=168), percentage of abnormal mitochondria in different groups (% abnormal mitochondria) (n=6 for each group). \**P*<0.05; \*\**P*<0.01; \*\*\**P*<0.001. **C,** ATP content in hearts from Sham, TAC and TAC+Nrg2 group (n=3 for each group). \**P*<0.05; \*\**P*<0.01; ns, not significant. **D,** NRVMs were treated with vehicle, Ang II or Ang II and recombinant Nrg2 for 48 h, and then stained with Mitotracker Red for mitochondria and DAPI for nuclei. Scale bar=20 μm. **E,** Quantification of cell mitochondrial fission rate (n=3). \*\**P*<0.01; \*\*\**P*<0.001. **F,** NRVMs were treated with vehicle, Ang II or Ang II and recombinant Nrg2 for 48 h, respectively. Oxygen consumption rate (OCR) of NRVMs was measured using a Seahorse flux analyzer (n=4). **G,** Analysis of OCR relating to basal respiration, ATP production-coupled respiration and maximal respiration shown in F (n=4). \*\**P*<0.01; ns, not significant. **H,** NRVMs were incubated with control medium, conditioned medium from CD163^+^/Dab2^+^ macrophages, and conditioned medium from Nrg2-knockdown CD163^+^/Dab2^+^ macrophages in response to vehicle or Ang II for 48 h. OCR was measured using a Seahorse flux analyzer (n=3). **I,** Analysis of basal respiration, ATP production coupled respiration and maximal respiration shown in H (n=3). \**P*<0.05; \*\**P*<0.01; \*\*\**P*<0.001; ns, not significant. **J,** NRVMs were incubated with control medium, conditioned medium from CD163^+^/Dab2^+^ macrophages, and conditioned medium from Nrg2-knockdown CD163^+^/Dab2^+^ macrophages in response to vehicle or Ang II for 48 h. Cardiomyocytes were stained with Mitotracker Red for mitochondria and DAPI for nuclei. Scale bar=20 μm. **K,** Quantification of the cell rate of mitochondrial fission (n=4). \*\**P*<0.01; \*\*\**P*<0.001; ns, not significant.

## Discussion

Pathological cardiac hypertrophy is closely associated with an increased risk of chronic heart failure^33^. Although inhibiting pathological hypertrophy is recognized as a crucial therapeutic strategy for heart failure, there is still a lack of effective targets that have been successfully translated into clinical practice^3^. Here, we identify a novel macrophage subset marked by high expression of CD163 and Dab2, and demonstrate its cardioprotective role during pathological cardiac hypertrophy. These CD163^+^/Dab2^+^ macrophages interact with cardiomyocytes via the Nrg2/ErbB4 signaling pathway. Conditioned medium derived from CD163^+^/Dab2^+^ macrophages alleviates cardiomyocyte hypertrophy; however, conditioned medium from Nrg2-knockdown CD163^+^/Dab2^+^ macrophages fails to confer protection against cardiac hypertrophy. Mechanistically, CD163^+^/Dab2^+^ macrophages restore mitochondrial homeostasis in cardiomyocytes through the Nrg2/ErbB4 pathway, thereby attenuating pathological cardiac hypertrophy. Thus, this study expands our understanding of macrophage-cardiomyocyte crosstalk and the function of Nrg2 during cardiac remodeling and suggests novel therapeutic strategies for pathological cardiac hypertrophy by targeting mitochondrial dysfunction (Figure S8).

The heterogeneity of cardiac macrophages and their distinct roles in injury and repair have been increasingly recognized^34^. These cells play critical roles in myocardial survival and adaptive cardiac remodeling, and are involved in the regulation of various pathological processes, including immune-inflammatory responses, energy metabolism, fibrosis, angiogenesis, and cardiomyocyte apoptosis^13,35^. In this study, we identify a distinct subpopulation of cardiac macrophages that exhibit relatively high expression of M2 macrophage markers CD163 and F13A, as well as the tumor suppressor Dab2. Disabled-2 (Dab2) is a clathrin- and cargo-binding endocytic adaptor protein with diverse functions in multiple signaling pathways, including the regulation of cellular differentiation, proliferation, migration, tumor suppression, and other essential homeostatic processes^36^. Notably, Dab2 has been identified as a putative regulator of cardiomyocyte development, promoting cardiomyocyte differentiation through the inhibition of Wnt/β-catenin signaling^37^. Additionally, Dab2 plays a critical role in macrophages, where its overexpression of Dab2 suppresses the pro-inflammatory polarization of macrophages induced by HDAC5^38^. Dab2 is upregulated in M2 macrophages, where it inhibits the pro-inflammatory M1 phenotype and modulates inflammatory signaling during macrophage polarization^27^. Furthermore, the inhibition of M1 polarization in macrophages by remimazolam has been shown to mitigate cardiac ischemia-reperfusion injury^39^. Similarly, latifolin protects the heart from doxorubicin (DOX)-induced cardiotoxicity by reducing the ratio of M1 to M2 polarized macrophages^40^. Therefore, these results suggest that Dab2 may regulate cardiac function through modulation of macrophage phenotypic switching. Here, we promulgate that CD163^+^/Dab2^+^ macrophages exert negative regulation on cardiac hypertrophy via the Nrg2/ErbB4 signaling pathway. However, the interplay between Dab2^+^ TAMs (tumor associated macrophages) and FAP^+^ CAFs (cancer-associated fibroblasts) has been shown to contribute to the formation of an immunosuppressive barrier and resistance to immunotherapy in hepatocellular carcinoma (HCC)^41^. Collectively, these findings indicate that distinct Dab2^+^ macrophage subsets can perform diverse functional roles across different microenvironments and disease states, underscoring their remarkable plasticity and adaptability in immune responses.

Crosstalk between cardiomyocytes and non-cardiomyocytes plays a crucial role in the progression of pathological cardiac hypertrophy^42,43^. In the present study, we demonstrate that CD163^+^/Dab2^+^ macrophages interact closely with cardiomyocytes through the ligand-receptor pair Nrg2/ErbB4. Nrg2 expression level is positively correlated with the presence of CD163^+^/Dab2^+^ macrophages. However, the interaction between CD163⁺/Dab2⁺ macrophages and cardiomyocytes via the Nrg2/ErbB4 signaling pathway is diminished during pathological cardiac hypertrophy, likely due to reduced expression of the ligand-receptor pair Nrg2/ErbB4. Previous study has shown that the neuregulin receptors ErbB2 and ErbB4 are downregulated at the stage of early heart failure in rats of chronic hypertrophy secondary to aortic stenosis^44^. Importantly, neuregulins (Nrg1 and Nrg2) and their receptors (ErbB2 and ErbB4) are essential for cardiac development, as they regulate myocardial growth and promote the survival of ventricular cardiomyocytes in embryonic, postnatal, and adult rats^45,46^. Nrg1 promotes cardiomyocyte migration and cell cycle progression during ventricular development^47^, positioning it as a promising therapeutic target for cardiac repair^48^. Our study demonstrates that administration of recombinant Nrg2 attenuates pathological cardiac hypertrophy by restoring cardiac function, and reducing cardiac fibrosis and apoptosis. In contrast, blocking the Nrg2/ErbB4 signaling pathway eliminates these cardiac protective effects, suggesting that Nrg2 may serve as a potential therapeutic target for pathological cardiac hypertrophy.

The pathophysiological progression of cardiac hypertrophy involves cardiac metabolic remodeling, characterized by the compensatory activation of glycolytic pathways, impaired mitochondrial biogenesis, and diminished oxidative phosphorylation capacity^33,49^. Our multi-omics data unveil the elevated glycolysis activity and depressed TCA and FAO activities, during pathological cardiac hypertrophy. Metabolic remodeling, together with mitochondrial dysfunction, contributes to energy deprivation in the failing heart^50^. Mitochondrial dysfunction plays a pivotal role in the progression of pressure overload-induced cardiac hypertrophy and is closely linked to increased morbidity and mortality^11,51,52^. Interestingly, CD163^+^/Dab2^+^ macrophages ameliorate Ang II-induced mitochondrial dysfunction through the Nrg2/ErbB4 signaling pathway. Jiayu Ren et al have demonstrated that macrophage-specific SGK3 deficiency exacerbates Ang II-induced myocardial fibrosis and mitochondrial oxidative stress^53^. Collectively, these findings highlight the regulatory role of macrophages in modulating mitochondrial function in cardiomyocytes. Mitochondrial function is increasingly recognized as a potential therapeutic target for heart failure, aimed at directly improving cardiac performance^17^. In our present study, we show that administration of recombinant Nrg2 to TAC mice at an early stage of myocardial hypertrophy restores mitochondrial bioenergetics and thereby attenuates pathological cardiac hypertrophy. Indeed, although further detailed investigations are required to fully elucidate the mechanisms by which CD163^+^/Dab2^+^ macrophages orchestrate cardiac hypertrophy and mitochondrial function via the Nrg2/ErbB4 pathway, our findings suggest that this axis represents a promising therapeutic strategy for the treatment of pathological cardiac hypertrophy.

In summary, we have gathered evidence that CD163^+^/Dab2^+^ macrophages communicate with cardiomyocytes via the Nrg2/ErbB4 pathway to improve mitochondrial bioenergetics, thereby alleviating cardiac hypertrophy. Conceptually, our findings highlight the role of the Nrg2/ErbB4 pathway in cardiac remodeling, suggesting it may serve as a potential therapeutic target for the treatment of pathological cardiac hypertrophy.

## Acknowledgments

The authors would like to thank the Proteomics and Metabolomics Platform (Guangzhou Laboratory) for liquid chromatography-time-of-flight mass spectrometry (Q-TOF), and Dr. Nannan Wang for raw data collection and analysis.

## Footnotes

## Nonstandard Abbreviations and Acronyms

TAC: transverse aortic constriction
NRVM: neonatal rat ventricular myocytes
BMDM: Bone marrow-derived macrophages
Anp: atrial natriuretic polypeptide
Bnp: brain natriuretic peptide
ETC: electron transport chain
OCR: oxygen consumption rate
Myh7: Myosin heavy chain 7
Col1a1: Collagen type I alpha 1 chain
Col3a1: Collagen type III alpha 1 chain
Pfkp: Phosphofructokinase
Hk2: Hexokinase 2
Bpgm: Bisphosphoglycerate mutase
Aldh2: Aldehyde dehydrogenase 2
Fbp2: Fructose-bisphosphatase 2
MDH1: Malate dehydrogenase 1
Dld: Dihydrolipoamide dehydrogenase
Sdhb: Succinate dehydrogenase complex subunit B
Sdha: Succinate dehydrogenase complex subunit A
Pdha1: Pyruvate dehydrogenase E1 subunit α1
Acadm: Medium-chain acyl-CoA dehydrogenase
Acaa2: Acetyl-CoA acyltransferase 2
Decr1: 2,4-Dienoyl-CoA reductase 1
Glc: Glucose
GAP: Glyceraldehyde-3-phosphate
FBP: Fructose-1,6-bisphosphatase
WT: wild type
WGA: Wheat germ agglutinin

